# Uncertainty-guided learning with scaled prediction errors in the basal ganglia

**DOI:** 10.1101/2022.01.10.475599

**Authors:** Moritz Moeller, Sanjay Manohar, Rafal Bogacz

**Affiliations:** Nuffield Department of Clinical Neurosciences, University of Oxford; Department of Experimental Psychology, University of Oxford

## Abstract

To accurately predict rewards associated with states or actions, the variability of observations has to be taken into account. In particular, when the observations are noisy, the individual rewards should have less influence on tracking of average reward, and the estimate of the mean reward should be updated to a smaller extent after each observation. However, it is not known how the magnitude of the observation noise might be tracked and used to control prediction updates in the brain reward system. Here, we introduce a new model that uses simple, tractable learning rules that track the mean and standard deviation of reward, and leverages prediction errors scaled by uncertainty as the central feedback signal. We provide a normative analysis, comparing the performance of the new model with that of conventional models in a value tracking task. We find that the new model has an advantage over conventional models when tested across various levels of observation noise. Further, we propose a possible biological implementation of the model in the basal ganglia circuit. The scaled prediction error feedback signal is consistent with experimental findings concerning dopamine prediction error scaling relative to reward magnitude, and the update rules are found to be consistent with many features of striatal plasticity. Our results span across the levels of implementation, algorithm, and computation, and might have important implications for understanding the dopaminergic prediction error signal and its relation to adaptive and effective learning.

**Author Summary:** The basal ganglia system is a collection of subcortical nuclei in the mammalian brain. This system and its dopaminergic inputs are associated with learning from rewards. Here, dopamine is thought to signal errors in reward prediction. The structure and function of the basal ganglia system are not fully understood yet—for example, the basal ganglia are split into two antagonistic pathways, but the reason for this split and the role of the two pathways are unknown. Further, it has been found that under some circumstances, rewards of different sizes lead to dopamine responses of similar size, which cannot be explained with the reward prediction error theory. Here, we propose a new model of learning in the basal ganglia—the scaled prediction error model. According to our model, both reward average and reward uncertainty are tracked and represented in the two basal ganglia pathways. The learned reward uncertainty is then used to scale dopaminergic reward prediction errors, which effectively renders learning adaptive to reward noise. We show that such learning is more robust than learning from unscaled prediction errors and that it explains several physiological features of the basal ganglia system.

## Introduction

For any organism, better decisions result in better chances of survival. Reward prediction is an important aspect of this—for example, if an organism can predict the size of a food reward associated with some behavior, it can decide whether it is worth to engage in that behavior or not. Reward predictions are typically based on values learned from previous reward observations. An extensive literature describes the role of reward prediction in behavior, as well as the related neural mechanisms (1).

Piray and Daw (2) argue that when trying to predict rewards, the organism faces two challenges. The first challenge is the dynamic nature of the environment: reward sizes and contingencies might change over time, in ways that cannot be predicted. Such genuine changes in the environment can be quantified by the typical rate of change, which is called ***process noise***. The second challenge is ***observation noise***: even if the environment is stable, rewards will vary from experience to experience. This could be due to the random nature of the environment, but also to variability in the organism’s own behavior, or to noise in the organism’s perception and evaluation systems.

The stock market serves as a nice example of the two types of noise: consider the day-to-day change of a stock price as a reward signal (if the stock price rises from 20 GBP to 21 GBP overnight, then the shareholders win 1 GBP in that transition). Most of the variability of that signal will be due to random fluctuations—this can be classified as observation noise. However, a part of the signal’s variability will reflect genuine lasting changes in the stock prize, for example a rise in price when a new product is released. This part should be classified as process noise.

What is the best reward prediction method an organism could use when facing process noise and observation noise? Similar problems occur in engineering, for example in the context of navigation. There, a very versatile solution has been found. That solution, called the ***Kalman filter*** (3), is very widely used—it even played a role in the moon landing (4). The Kalman filter describes how estimates of a variable must be updated when new noisy observations of that variable become available. For certain types of signals, it can be shown that the Kalman filter is indeed the ***optimal*** method for prediction in the presence of noise. The method has proven useful not only in engineering, but also as a model of neural and behavioral processes (5-8).

However, if one wants to use a Kalman filter to predict rewards, one runs into a problem: the Kalman filter requires estimates of the magnitudes of both process noise and observation noise as parameters. Where to take these values from? An organism might either use fixed (perhaps genetically determined) values or estimate the values somehow. The former option bears a risk: if the world changes, the quality of the organism’s predictions might decline strongly. The latter option raises the next question: how is this estimation done?

Solutions for this have been proposed. For example, Piray and Daw (9) present a model that tracks both process noise and observation noise alongside reward, allowing for ***adaptive*** Kalman filtering. However, their model (a variant of the particle filter) is targeted at the computational level, i.e., it is set up to investigate how the simultaneous adaptation to two noise types of noise affects learning. Questions concerning the underlying biological mechanisms remain largely unaddressed. The model of Piray and Daw (9) is hence not suitable to describe biological learning on the mechanistic or the algorithmic level.

This leads us to the central question of this paper: ***how might organisms track observation noise in a biologically plausible, computationally simple way, and use it for adaptive reward prediction?*** We propose that observation noise is tracked in the basal ganglia, and that it is used to improve learning performance by normalizing reward prediction errors. This proposal is based on the observation of dopamine activity patterns consistent with normalized prediction errors (10), as well as on previous suggestions that reward uncertainty might be represented in the basal ganglia (11, 12). Such scaling of prediction errors is conceptually related to scaling mechanisms that occur in free energy models, as well as to techniques such as adaptive momentum that are used to improve fitting algorithms (see Discussion for details).

Below, we give a detailed analysis of how the basal ganglia circuit might carry out the computations necessary to track and utilize reward observation noise. To provide some context for this analysis, we now move on to a brief review of the main features of that part of the brain.

Fig 1 shows a highly simplified version of the cortico-basal-ganglia-thalamic circuit, with three important brain regions—the cortex, the striatum, and the thalamus—arranged along the vertical axis.

**Fig 1.**
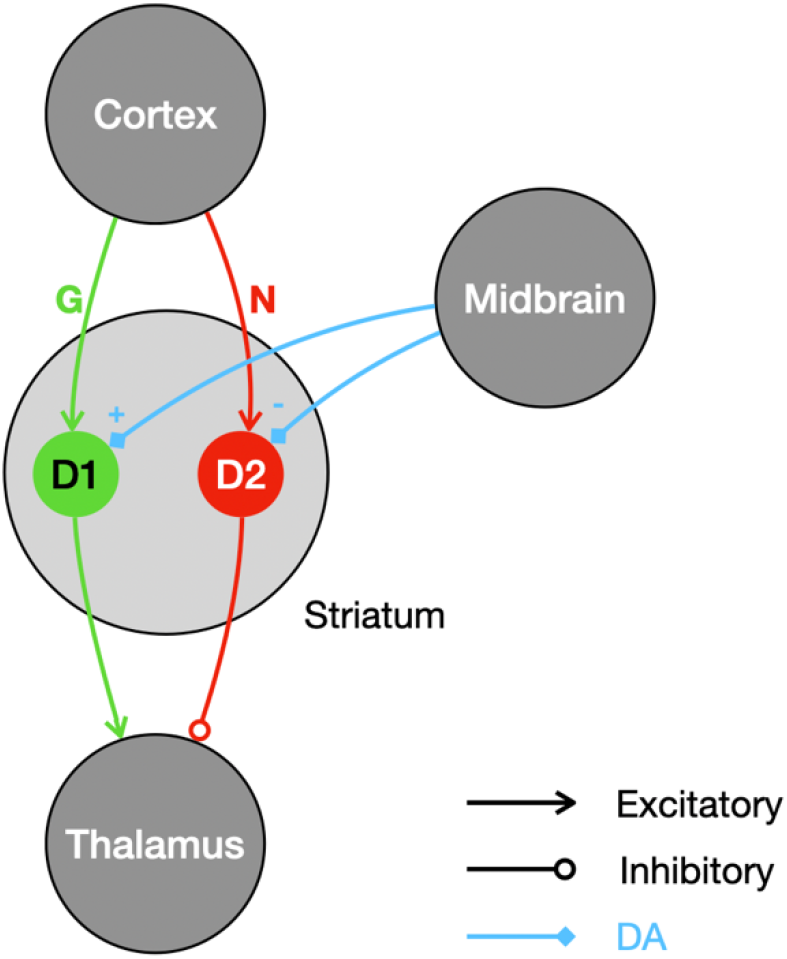
The simplified basal ganglia circuit. Selected nuclei and connections are shown as circles and arrows. Green connections correspond to the direct pathway; red connections correspond to the indirect pathway. Dopamine projections are shown in blue.

The striatum is the largest nucleus within the basal ganglia system. It includes two populations of medium spiny projection neurons (SPNs): the D1 and the D2 population (D1 and D2 are the types of dopamine receptors that the corresponding neurons express). This division of the striatum gives rise to two parallel descending pathways called the direct/Go and the indirect/No-go pathway, shown in green and red respectively in Fig 1 (13-15). The cortical inputs to these two pathways are modulated by the strengths of the synapses between cortex and the striatal populations, collectively labeled *G* and *N* in Fig 1.

The thalamus receives the output of the basal ganglia circuit. The effects of the direct and the indirect pathway on the thalamus are differential: the direct pathway effectively excites the thalamus; the indirect pathway effectively inhibits it. Note that the projections from the striatum to the thalamus in Fig 1 are abstractions—in the brain, there are several intermediate nuclei between the striatum and the thalamus.

The final key element of the basal ganglia system are the dopamine projections from midbrain regions that target the striatal D1 and D2 populations. The effects of dopamine release on the striatal populations are twofold: dopamine modulates activity, but also triggers plasticity. The direction of those effects depends on the receptor type of the target neuron: dopamine increases excitability and potentiates synapses in the D1 population while it decreases excitability and depresses synapses in the D2 population (13-15).

Overall, we may view the basal ganglia circuit as two parallel descending pathways that converge on the level of the thalamus, where they have opposite effects. Those pathways are differentially modulated by dopamine, which also controls synaptic plasticity between the cortex and the striatum.

Concerning the function of the elements of this model of the basal ganglia, we follow a popular view often used in modelling (11, 16, 17): the cortex supplies contextual information, i.e., cues, stimuli, sensory data or information on the state of the environment; the other populations (D1, D2 and Thalamus) encode actions. Each action is represented by a distinct subpopulation of each nucleus, and the connectivity between the nuclei is action specific. For example, assume there is a subpopulation in D1 associated with pressing a lever. A corresponding subpopulation could be found in D2 as well as in the thalamus, which is known to relay motor commands to the relevant cortical areas (18). The two striatal subpopulations associated with the lever press would then project exclusively to the lever-press subpopulation in the thalamus, together forming what is often called an action channel (19). Learning is assumed to take place at the interface between the cortex and the striatum (which, in this model, can be considered a state-action mapping). Learning is implemented through dopamine-mediated plasticity of cortico-striatal synapses. These synapses within an action channel are assumed to store information on the action value (the mean reward associated with the action). Action values determine the relative activations of action channels (i.e., the difference in activation between the Go- and the No-go pathways), and hence contribute to action selection at the level of the thalamus. It has been proposed (11) that cortico-striatal synapses additionally encode reward uncertainty (in the sum of the weights in the Go- and the No-go pathways), as we explain in detail below.

In the following sections, we present and analyze a model—the scaled prediction error (SPE) model—that tracks observation noise and uses it for adaptive reward prediction. In the first part of the paper, we introduce the model and test its performance. There, we show that it outperforms the classic Rescorla-Wagner (RW) model of associative learning (20) which does not adapt to observation noise, using simulations of a reward prediction task. In the second part of the paper, we discuss neural mechanisms that might implement the SPE model in the basal ganglia circuit. We first focus on dopamine signals, and then move on to the mechanisms behind tracking observation noise and scaling prediction error.

## Results

### The model

The SPE model is a model of reward prediction—it predicts the magnitude of the next reward based on previous reward observations. It can be understood as approximate Bayesian inference with respect to a particular model of the reward generation process. This model is given by

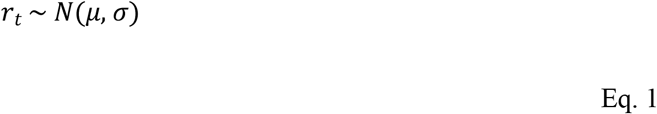

with *r*_*t*_ the reward in trial *t, μ* the mean reward and *σ* the reward observation noise. The SPE model can be derived by approximating Bayesian inference of the parameters *μ* and *σ* from the observed rewards (we show this derivation in Appendix S1). It does this by maintaining estimates *m* and *s* of those parameters, which it updates whenever a new reward is observed. Though the model is designed to infer the mean and standard deviation of stationary reward processes, it can also be applied to reward processes with a drifting mean (i.e., to processes with non-zero process noise). We show this below in our simulations.

The SPE update rules are

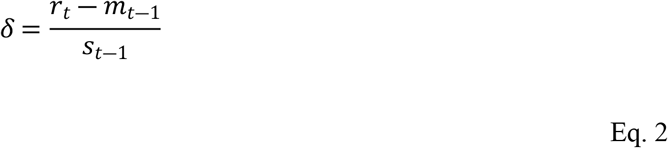

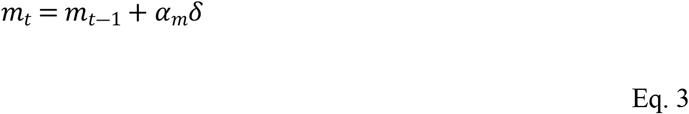

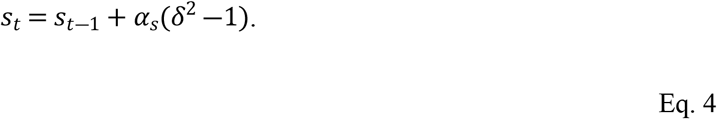

In these equations *δ* is a scaled reward prediction error, while *α*_*m*_ and *α*_*s*_ denote the learning rates for the mean and standard deviation, respectively. We will next show that the equations can indeed recover mean reward and its standard deviation.

### Fixed point analysis

Do the SPE rules do what they are meant to do? Here, we use a stochastic fixed-point analysis to show that in theory, *m* and *s* should converge to the mean and the standard deviation of the reward signal. Let us assume that rewards are indeed generated by sampling from a distribution with mean *μ* and standard deviation *σ* (this could be a normal distribution or any other distribution with well defined mean and standard deviation).

We consider a situation in which the learner has already found the correct values of the variables it maintains, i.e., *m* = *μ* and *σ* = *s*. From there, what are the **expected updates**? A straightforward calculation yields

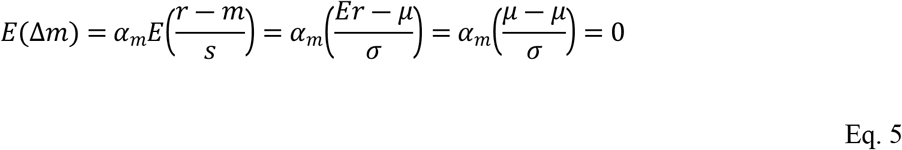

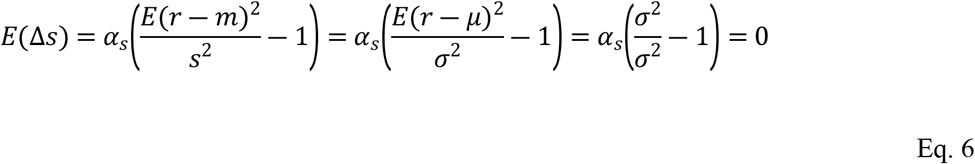

with *Er* = *μ* and *E*(*r* − *μ*)^2^ = *σ*^2^ by definition. We find that the expected change away from (*m, s*) = (*μ,σ*) is zero, which makes (*μ, σ*) a stochastic fixed-point. We may conclude that in equilibrium, the rules given in Eq. 2 – 4 should give us unbiased estimates of the reward mean and standard deviation.

It can be shown that the SPE model is related to the Kalman filter—it can be viewed as an approximation to a steady-state Kalman filter, which becomes more accurate if observation noise dominates process noise (see Appendix S2 for details). Finally, note that the RW model is a special case of the SPE model (i.e., if *α*_*S*_ =0).

### Performance

How do the SPE rules compare to established rules such as the Rescorla-Wagner model with respect to accurate reward predictions? To compare the performances of SPE learning and RW learning, we apply both to a reward prediction task: sequences of rewards are generated according to

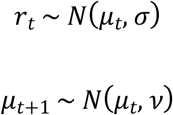

In the above equation *v* is the standard deviation of the process noise. Both learners observe the reward signal and provide reward predictions at every trial. The learners’ performance is judged by measuring the average precision of their predictions.

The task is designed to challenge the learners with rewards that change over time, forcing them to continuously learn. Note that the reward-generating process here is more complex than the generative model from which SPE learning is derived. This is not a problem—the SPE model is robust with respect to violation of its assumptions, as we shall see below. Of course, one could derive learning rules tailored to this reward process. These would involve a representation of the process noise, or volatility. This is not our goal here. Instead, we are interested in the SPE learning rules as a model of basal ganglia learning and want to test their performance.

We compare the models for different levels of observation noise *σ*, while keeping the process noise *v* constant at *v* =1. The results of those comparisons are presented in Fig 2 (see Methods for details of the implementation).

**Fig 2.**
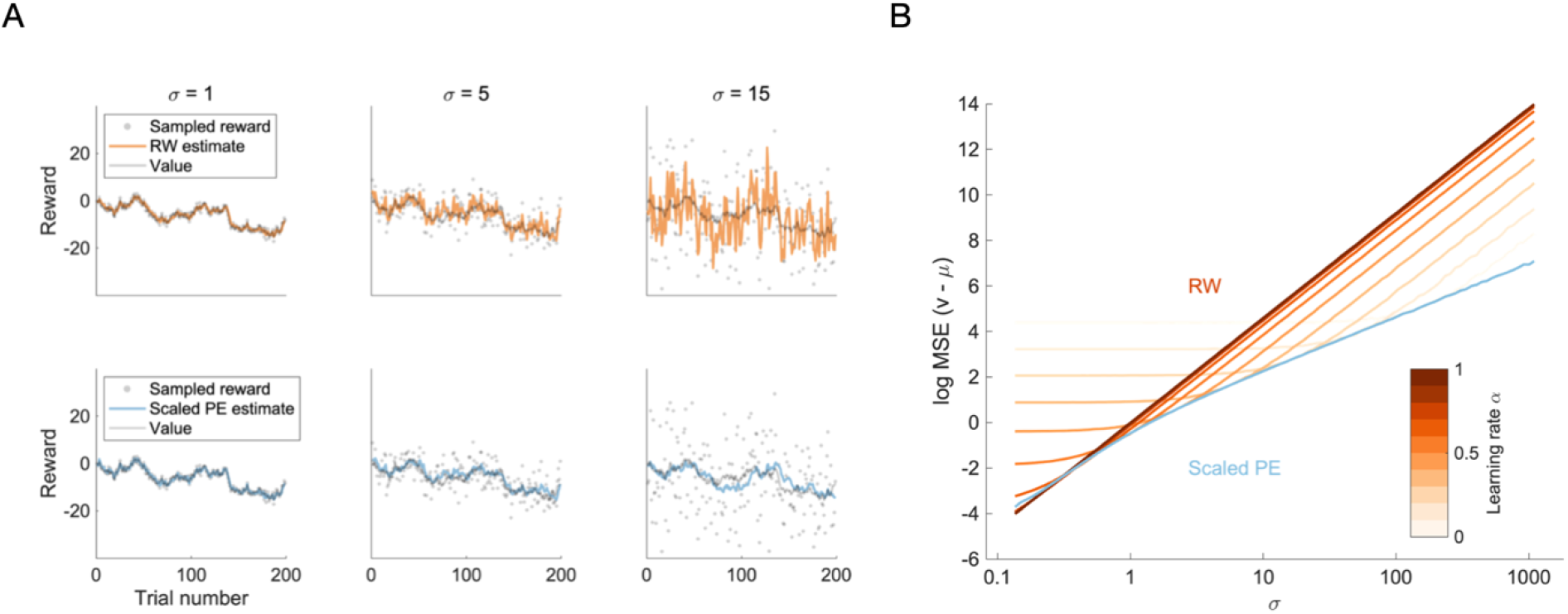
Reward prediction performance of the RW and SPE models. **A** The first 200 trials of reward prediction for the RW learner (upper row, orange color) and the SPE learner (lower row, blue color). The true value (grey line), the observed rewards (grey dots) and the learner’s estimate (colored line) are shown as a function of trial number. Columns correspond to selected levels of observation noise (σ = 1, 5, 15). **B** Learning performance averaged over trials. We show the logarithm of the mean squared difference between the mean of the reward distribution and the learner’s prediction thereof, as a function of the observation noise σ. Orange lines correspond to RW learners, the blue line corresponds to a SPE learner parametrized with α_m_ = 1 and α_S_ = 0.01. The different shades of orange correspond to different learning rates, as indicated by the color bar.

Looking first at the time series in Fig 2A, we find that there is a qualitative difference between the RW learner in the top row and the SPE learner in the bottom row: as the noise level *σ* increases, the RW learner’s predictions increasingly fluctuate, since the reward prediction errors (and hence the updates) scale proportionally to the amplitude of the observation noise. This is not so for the SPE learner, whose predictions fluctuate as much for low noise levels as they do for high noise levels.

This effect is also visible in the aggregated performance measure, shown in Fig 2B: the mean squared errors of the learners’ predictions grow with observation noise for all learners, but they grow stronger for the RW learners. We find a very stereotyped effect for the average performance of RW learners: as the level of noise increases, prediction accuracy does not change much up to a certain point (call this the plateau) and grows steadily after that point (call this the slope). This is the case irrespective of the learning rate. Smaller learning rates have a plateau that extends to higher noise levels but also provides a lower accuracy. The steepness of the slope is invariant across learning rates.

To gain an intuition for the shape of these curves, let us compare two different situations. First, consider very low levels of observation noise. In this regime, reward observations are almost identical to observation the underlying mean reward. Hence, if the observed reward changes, this mostly reflects a genuine change of the underlying mean reward. To keep reward predictions precise, such changes should be followed. However, for learning rates smaller than one, the RW model does not fully follow the changes of the reward signal—we may call this underfitting, as the model ignores meaningful variation in the signal. The resulting error dominates the performance. The magnitude of this error depends on the volatility of the signal. Since the volatility is kept constant in the simulations in Fig 2B, we see a performance plateau at low levels of observation noise.

Now, consider very high levels of observation noise. In this case, reward observations are very inaccurate—an observation tells us very little about the underlying mean reward. Changes in observed rewards mostly reflect the noisiness of the observations and should be ignored. However, for learning rates larger than zero, the RW model does not fully ignore those fluctuations, but follows them. This can be called overfitting, as the model tries to adapt to random fluctuations.

Overall, the behavior of the RW learners is such that for each given level of observation noise there is an optimal learning rate: if one selects any one position on the x-axis of Fig 2A, there is always a single orange curve with the lowest y-coordinate (and hence the smallest average error) at that position. In general, we find: the higher the observation noise, the lower the optimal learning rate. This appears consistent with intuition—if observation noise is high, there is less useful information in any single observation and an organism should therefore update its estimate more carefully.

The SPE learner shows different behavior. There is also a slope (prediction accuracy steadily decreases with increasing observation noise), but no plateau. The steepness of the slope changes at *σ* = 1. For higher levels of observation noise, the slope of the SPE learner is shallower than those of the RW learners. We find that for any given level of noise *σ* larger than one, the performance of the SPE model is about as good as the performance of the best RW model. This suggests that in the regime of high observation noise we might view the SPE model as an RW learner that reaches optimal performance by fine-tuning itself to the estimated level of observation noise.

Can one do better than this? In fact, one can show that the SPE model (parametrized with *α*_*m*_ = 1) is approximately optimal in the situation investigated here: for high levels of observation noise, SPE learning approximates the steady-state Kalman filter (we show this in Appendix S2), which is approximately optimal for the types of signals we use here.

As mentioned above, to use a Kalman filter, one needs to provide it with the correct values of *σ* and *v*. This is also true for the steady-state version of the Kalman filter, but it is not the case for the SPE model: here one only needs to provide *α*_*m*_ —which corresponds to *v* (see Appendix S2), —but not *σ*, which the model can track by itself. We can thus think of SPE learning as ***adaptive*** steady-state Kalman filtering.

However, to work optimally, the SPE model still needs to be provided with the correct value for *α*_*m*_. To make the model more autonomous, one might extend it with a mechanism to track *v* alongside *σ*, for example the mechanism proposed by Piray and Daw (2). This is an interesting direction for further research but goes beyond the scope of this work.

In summary, we find that the SPE model is approximately optimal for signals with *v* < *σ*. In particular, it will be at least as good as any RW learner, and about as good as a steady-state Kalman filter. SPE learners thus appear particularly well suited to track signals with unknown or changing levels of observation noise, as they can adapt themselves to whatever level of noise they experience. In contrast, an RW learner would either have to be fine-tuned based on prior knowledge, or it would perform suboptimally due to under- or overfitting.

### The neural implementation of SPE learning

Could the SPE learning rules be implemented in the dopamine system and the basal ganglia pathways? In this section, we propose a possible mechanism. We suggest that striatal dopamine release broadcasts scaled prediction errors, 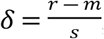, and that the update rules given in equations Eq. 3 and Eq. 4 are implemented by dopamine-dependent plasticity in the striatum. In the next subsections, we will analyze the plausibility of these suggestions. First, we discuss the relationship between dopamine responses and scaled prediction errors. Then we discuss how the SPE learning rules can be mapped on striatal plasticity rules. Finally, we propose a mechanism that might implement the scaling.

#### Scaled prediction errors are consistent with dopamine activity

In a seminal study, Tobler et al. (10) investigated how the responses of dopamine neurons to unpredictable rewards depended on reward magnitude, using electrophysiology in monkeys. Three different visual stimuli were paired with three different reward magnitudes (0.05 ml, 0.15 ml and 0.5 ml of juice). After being shown one of the stimuli, the monkeys received the corresponding reward with a probability of 50%. Seeing the stimulus allowed the monkey to predict the magnitude of the reward that could occur, but not whether it would occur on a given trial. Reward delivery thus came as a surprise and evoked a dopamine response. Interestingly, these responses did not scale with the magnitude of the received rewards. The measured dopamine responses are shown in Fig 3B.

**Fig 3.**
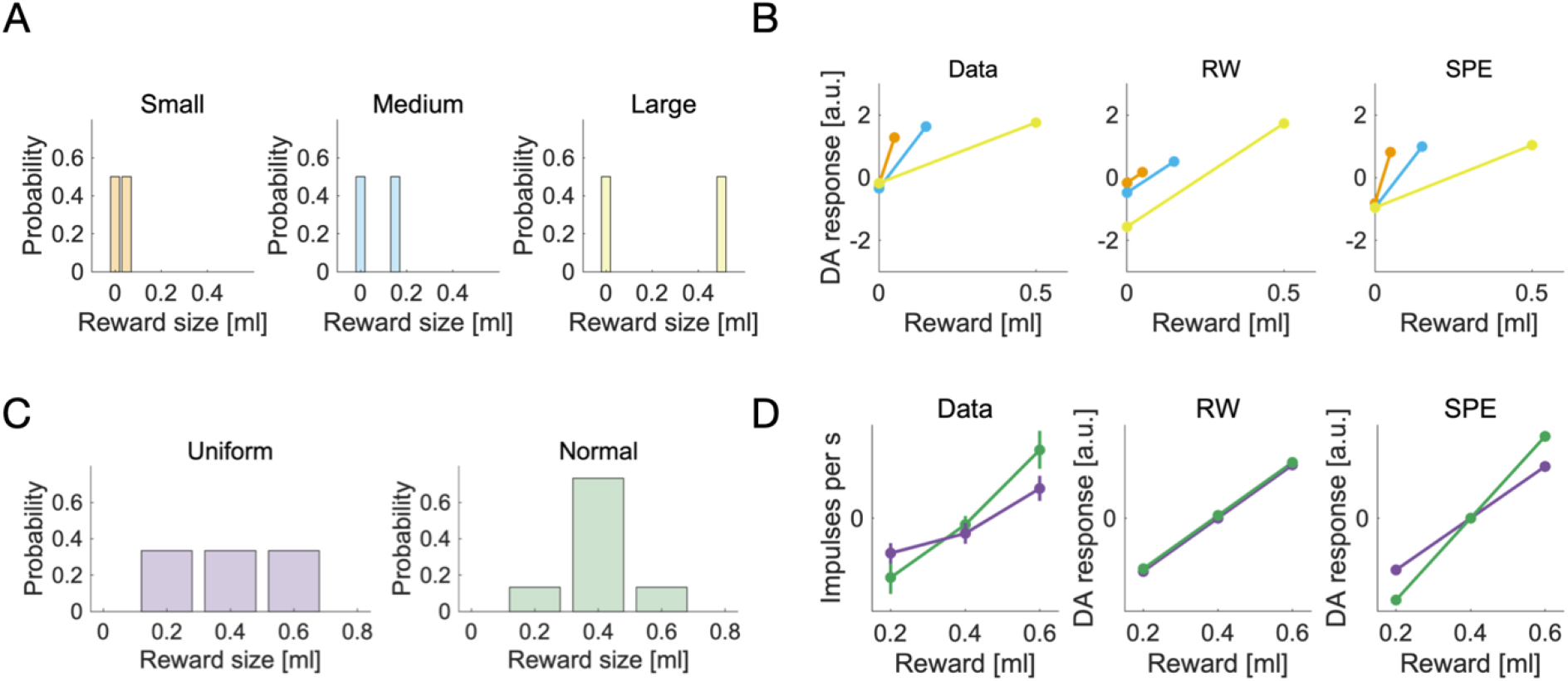
Dopamine responses to unpredictable rewards—experimental data and simulations. **A** The reward distributions used by Tobler et al. (10). Each distribution corresponds to an experimental condition. **B** Dopamine responses to rewards sampled from the distributions in A are shown as a function of reward magnitude, for the three different conditions. The representation of data is similar to that in figure 4C of Tobler et al. (10). We show experimental data, extracted from figure 4C (animal A) of Tobler et al. (10) and simulated data, using a standard RW model and the SPE model. The colors relate the dopamine responses in B to the reward distributions in A. **C** The reward distributions used by Rothenhoefer et al. (21). The panel is reproduced from Rothenhoefer et al. (21), figure 1A. **D** Dopamine responses to rewards sampled from the distributions in C. We show the empirical values, reproduced from Rothenhoefer et al. (21), figure 2E, and the responses according to the RW model computed analytically as δ = r − μ, and the SPE model computed as 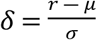, where µ and σ are the mean and standard deviation of corresponding reward distributions in C. Purple lines correspond to the uniform reward distribution, green lines correspond to the normal reward distribution.

This result was unexpected—standard RW learning would predict that the residual prediction errors in rewarded trials should grow linearly with reward magnitude. Our new SPE rules, on the other hand, predict exactly what has been observed. See Fig 3B for simulated and experimental DA responses. One may object that the results of Tobler et al. (10) might also be explained by scaling with respect to the reward range—reward range and reward standard deviation cannot be dissociated in that experiment. While that is true, another recent experiment can dissociate them: Rothenhoefer et al. (21) used two reward distributions with the same reward range but different reward standard deviations in a Pavlovian conditioning task (see Fig 3C).

After exhaustive training, single unit recordings were performed to measure dopamine responses to rewards that deviated from the expected value. It was found that the same deviation from the expected value caused stronger dopamine responses for the distribution with the smaller standard deviation (Fig 3D, first panel). This is consistent with scaling by reward standard deviation, but not with scaling by reward range---both distributions had the same range, so scaling by range should yield similar responses for both conditions. These experimental data cannot be accounted for by the RW model (Fig 3D, second panel), but can be reproduced by the SPE model (Fig 3D, third panel).

#### The SPE learning rules are consistent with striatal plasticity

After establishing that dopaminergic scaled prediction errors are plausible, we now move on to discuss how the update rules given in Eq. 3 and Eq. 4 could be implemented in the basal ganglia circuit.

Mikhael and Bogacz (11) proposed a distributed encoding of the two reward statistics (*m* and *s*) in the two main basal ganglia pathways: in their model, the mean of the reward signal is encoded in the difference between synaptic inputs to striatal neurons in direct (Go) and indirect (NoGo) pathways, whereas the standard deviation of the signal is encoded in the sum of these inputs. Formally, we write

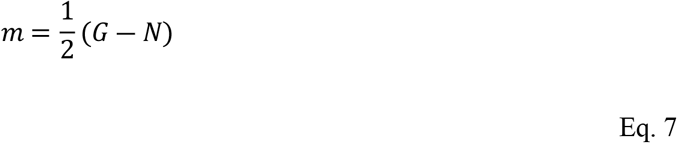

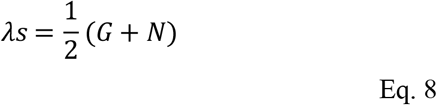

In Eq. 7 and 8, *G* and *N* denote the synaptic inputs in the direct and indirect pathway respectively (22), and *λ* is a coefficient determining the accuracy with which the standard deviation can be encoded (as explained below). These assumptions can be used to rewrite the learning rules given in Eq. 3 and 4 in terms of *G* and *N*. In particular, note that by combining Eq. 7 and 8, we see that *G* = *m* + *λs* and *N* = *λs* − *m*. Therefore, we can derive the update rules for *G*(or *N*) by adding (or subtracting) Eq. 3 and 4, and obtain

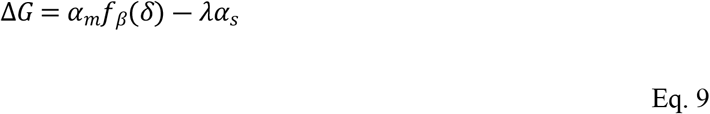

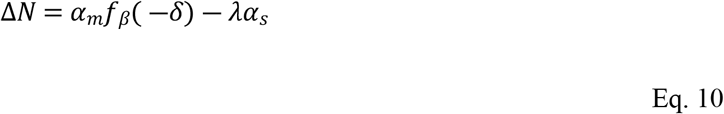

with *f*_*β*_(*δ*) = *βλδ*^2^ +*δ* and *β* = *α*_*s*_/*α*_*m*_. It is worth emphasizing that Eq. 9 and 10 are equivalent to Eq. 3 and 4 (because they are just rewritten in terms of different variables). Therefore, the model described by Eq. 9 and 10 estimates exactly the same mean and variances as a model described by Eq. 3 and 4, and hence it produces identical performance in Fig 2 and dopaminergic responses in Fig 3.

One important issue while considering biological plausibility of the model is the fact that the synaptic weights on the indirect pathway *N* cannot be negative, while the model assumes that these weights encode *N* = *λs* − *m*. Imposing a constraint of *N* being non-negative will limit the ability of the network to accurately estimate standard deviation of rewards to cases when it is sufficiently high (i.e. *σ* ≥ *μ/λ*). Hence the parameter *λ* controls the accuracy with which the standard deviation can be estimated. However, according to Eq. 8 there is a cost of high accuracy, because a high value of *λ* will result in overall larger values of the synaptic weights (analogously as in the model of Mikhael and Bogacz (11)), and hence higher metabolic cost of the computations.

Eq. 9 and 10 show three main features: 1) different overall effects of dopamine on plasticity in each pathway, 2) nonlinear effects of dopaminergic prediction errors represented by the transformations *f*_*β*_ and 3) synaptic unlearning represented by decay terms. We will discuss the experimental data supporting the presence of these features in turn.

First, the efficacy of direct pathway synapses is assumed to increase as a result of positive reward prediction errors (i.e., *δ* > 0), and decrease as a result of negative reward prediction errors (i.e., *δ* < 0). The opposite is assumed to hold for indirect pathway synapses: their efficacy should decrease with positive prediction errors and increase with negative prediction errors. This premise corresponds to the sign of the prediction error in Eq. 9 and 10, and it is consistent with data obtained in experiments (23).

Second, it is assumed that for striatal neurons in the direct pathway, positive prediction errors have a stronger effect on plasticity than negative prediction errors. This assumption is expressed in the shape of the function *f*_*β*_(*δ*), which is plotted in Figure 4Ai. Note that the slope for positive *δ* is steeper than for negative *δ*, implying that positive prediction errors should lead to bigger changes in *G* than negative prediction errors. The computational role of this nonlinearity is to filter the reward prediction errors: it amplifies the positive components while dampening the negative components of the signal.

**Fig 4.**
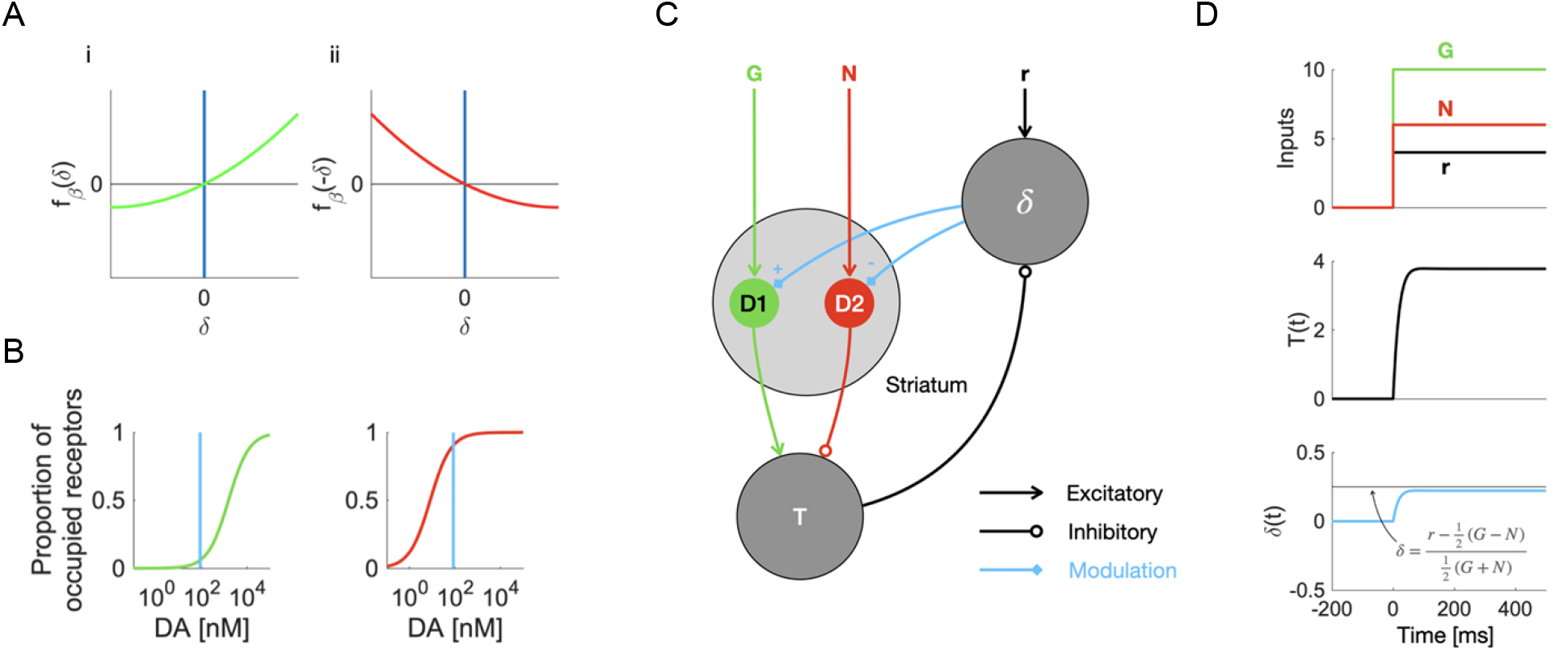
Plasticity and computations in the basal ganglia circuit. **A** The nonlinear transformation of dopaminergic prediction errors in the SPE model. The transformation in the direct pathway (i) and the transformation in the undirect pathway (ii) are mirror images of each other. **B** We plot the proportion of occupied receptors in the striatum as a function of dopamine concentration. The curves are based on the results of Dreyer et al. (25). The blue vertical lines indicate the baseline dopamine concentration in the ventral striatum, based on the results of Dodson et al. (26). The green curve corresponds to the occupancy of D1 receptors, the red curve corresponds to the occupancy of D2 receptors. Panel B is a partial reproduction of figure 3D of Möller and Bogacz (27). **C** The connectivity underlying a dynamical model of the simplified basal ganglia circuit. Circles correspond to neural populations; arrows between them indicate connections. **D** The computation of a scaled prediction error in continuous time, according to a dynamical model of the basal ganglia. We show how the relevant variables, T and δ, evolve as a function of time, assuming a step-function activation for the input nodes G, N and r. The black line in the lowest panel indicates the level of dopamine required for exact SPE learning.

For striatal neurons in the indirect pathway, the SPE model assumes the opposite: negative prediction errors should have a stronger plasticity effect than positive prediction errors, because the weight modification is proportional to *f*_*β*_(−*δ*), which is plotted in Figure 4Aii. Mikhael and Bogacz (11) argue that this premise is realistic, based on the different affinities of the D1 and D2 receptors that are present in striatal neurons in the direct and indirect pathways respectively: while D1 receptors are mostly unoccupied at baseline dopamine levels, D2 receptors are almost saturated—this is visualized in Fig 4B. Due to this baseline setting, additional dopamine should lead to a large difference in the occupation of D1 receptors and hence affect the neurons on the direct pathway, but only a small change in the occupancy of D2 receptors thus little influence the neurons on the indirect pathway. A decrease in dopamine, on the other hand, is strongly felt in D2 receptor occupancy but does not change D1 receptor occupancy much.

Third, an activity-dependent decay (or ‘unlearning’) is assumed to occur in the synaptic weights whenever they are activated in the absence of prediction errors. This is reflected in terms −*λα*_*s*_ in Eq. 9 and 10. On the neural level, that premise translates into mild long-term depression after co-activation of the pre- and postsynaptic cells at baseline dopamine levels. Recently, this effect has been observed at cortico-striatal synapses in vivo (24): in anaesthetized rats, presynaptic activity followed by postsynaptic activity caused LTD at baseline dopamine levels (i.e. in the absence of dopamine-evoking stimuli).

In summary, we discussed the three premises of the learning rules—the different overall effects of dopamine on plasticity in each pathway, the nonlinear effects of dopaminergic prediction errors and synaptic unlearning. We saw that all three premises are supported by the physiological properties of striatal neurons on the direct and indirect pathway.

#### Scaled prediction errors and the basal ganglia circuit

Next, we discuss how the scaled prediction error 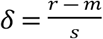 might be computed in the basal ganglia system. Expressed in terms of *G* and *N*, the scaled prediction error is given as

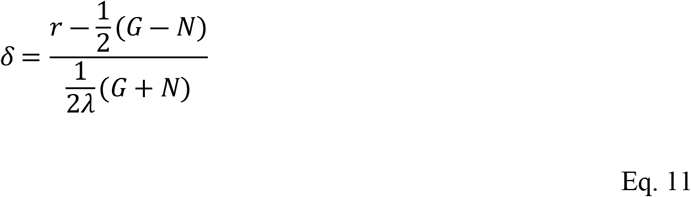

This seems to be a complicated combination of terms, and it is difficult to see how a simple network might compute it. Surprisingly, there is a simple approximate implementation based on a feedback loop. Here, we will describe that mechanism, using a minimal dynamical model of the basal ganglia network.

First, where is the feedback loop? Let us assume that the prediction error *δ* is encoded in the activity of a population of dopaminergic neurons. This population receives inhibitory input *T* from the thalamus, and excitatory input *r* that encodes a reward signal. Formally, we assume *δ* = *r* − *T*. We follow Möller and Bogacz (27) in assuming that the thalamic activity reflects the total output from the basal ganglia—the difference between the activity in the direct and indirect pathways—which is captured by

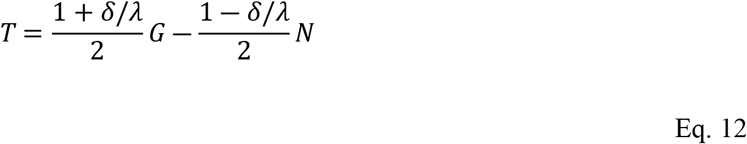

The first term 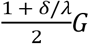 corresponds to the activity in the direct pathway, which is proportional to synaptic input *G*, and is increased by the dopaminergic modulation, because the gain of striatal neurons in the direct pathway is enhanced by dopamine. The second term 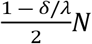 corresponds to the activity in the indirect pathway, which is attenuated by dopamine, because the gain of striatal neurons in the indirect pathway is reduced by dopamine (13, 14). The proposed model contains a feedback loop: dopamine release modulates the thalamic activity, which itself inhibits dopamine release.

To examine the computation of scaled prediction errors, we model the relevant populations’ activities as leaky integrators with effective connectivity as sketched in Fig 4C, using differential equations in continuous time.

The dynamical system sketched in Fig 4C corresponds to a set of differential equations,

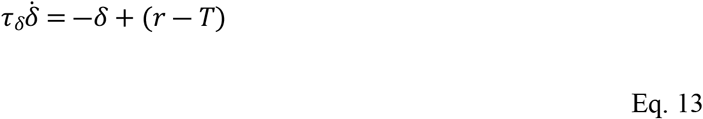

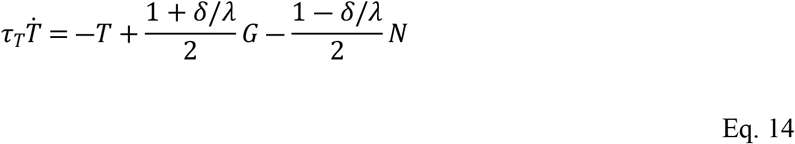

Here, *τ*_*δ*_ and *τ*_*T*_ are the characteristic timescales of the striatal dopamine release and thalamic activation. The system is set up such that its equilibrium point is consistent with our trial-wise description (*δ* = *r* − *T* and 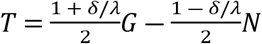 at 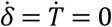). This asserts that the two levels of description are consistent with each other. Using these equilibrium equations, we can determine the equilibrium value of *δ* (by inserting one equation into the other and solving for *δ*). We find

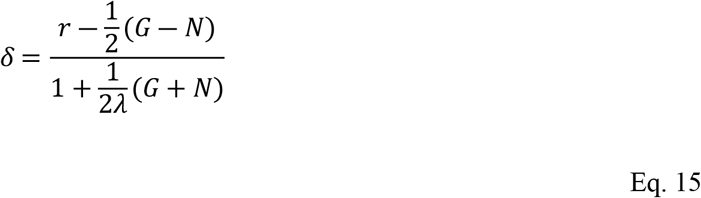

For 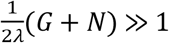, this approximates the scaled prediction error in Eq. 11. This suggests that the circuit can compute an approximation to the scaled prediction error. The approximation will be accurate as long as *s* is sufficiently large (recall from Eq. 8 that 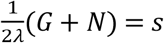). Although the additional term 1 in the denominator prevents perfect scaling, it might in fact be beneficial: it could prevent catastrophically large prediction errors that might cause the instabilities when the denominator becomes very small.

So far, it looks as though the circuit has an equilibrium point at approximately the right value. However, it is not yet clear whether and how this equilibrium is reached. To learn more about these aspects, we need to simulate the system. To simulate the computation of the prediction error, we assume *G, N* and *r* to be provided externally, for example through cortical inputs. *G* and *N* then represent precisely timed reward predictions, while *r* represents the reward signal itself. We model *G, N* and *r* as step-functions that jump from zero to their respective values at the same time, as illustrated by the first panel of Fig 4D. The time constants *τ*_*δ*_ and *τ*_*T*_ are set to realistic values taken from the literature (see Methods for details). A simulation of the system is shown in Fig 4D. We find that *δ* settles to its equilibrium value quite quickly (after tens of milliseconds) and without oscillations. This is likely due to the difference in time constants—the thalamic activity changes much faster than the striatal dopamine concentration. Our results suggests that even a simple system as the one in Fig 4C can compute scaled prediction errors through a feedback loop.

## Discussion

Above, we presented a new model of error-driven learning: the SPE model. We tested it in simulations and compared it with neural data. Now, we will discuss the new model more broadly. First, we will summarize our key findings. Then, we will present several testable predictions that follow from the model. Finally, we will discuss how the SPE model relates to other models from neuroscience and from artificial intelligence.

### Summary

This work introduces the SPE model, which describes how an organism might adapt its learning mechanism to changing levels of reward observation noise *σ*. First, we proposed the SPE learning rules, which can track the mean and standard deviation of a reward signal. We then tested the performance of the new rules. Comparing SPE learning with RW learning, we found that the new learning rules can improve performance when a learner faces unknown or varying levels of reward observation noise. Next, we reviewed empirical evidence relating to SPE learning. On the neural level, we found that SPE learning describes dopamine responses better than conventional models in several studies. We further showed how the basal ganglia pathways might implement the learning rules of the SPE model, and how scaled prediction errors could be computed in a dopaminergic feedback loop.

### Experimental predictions

Our model makes several predictions on different levels of analysis. First, SPE learning can be distinguished from other types of learning on the level of behavior. This is because according to SPE, the learning rate (and hence the speed of learning) should depend on the stochasticity of the reinforcement that drives learning. From this, different predictions follow.

For example, reward magnitude should ***not*** affect instrumental learning speed if animals were exposed to the reward prior to the instrumental phase. This is because SPE learning would allow the animal to normalize the reward magnitude through the adaptive scaling of prediction errors, as in the experiment by Tobler et al. (10). The dopamine-guided instrumental learning through normalized prediction errors would then not be affected by the overall scale of the rewards. Concretely, this could be tested in a decision-making task with two conditions. In one condition, correct choices are reinforced with high rewards (for example 0.6 ml of juice). In the other condition, low rewards are provided, for example 0.2 ml of juice. Conditions must be cued, perhaps by two different visual stimuli that precede the decision. According to the RW model, we expect to see a steeper learning curve in the high reward condition, as in the experiment of (28). However, SPE theory predicts that this difference between conditions should vanish if an appropriate pretraining is applied. For example, the cues could be associated with the different reward sizes through Pavlovian conditioning. According to SPE theory, the pretraining should establish a condition-specific normalization cued by the stimuli. This normalization should then lead to normalized learning curves in the instrumental phase. Ultimately, the difference between the learning curves should vanish.

Furthermore, SPE learning predicts that learning rates should change if reward stochasticity is changed. This could be tested by having participants track and predict a drifting reward signal which shows different levels of stochasticity at different times. If participants use SPE, the learning rate should decrease with increasing stochasticity, which would lead to invariant update magnitudes. On the other hand, if participants do not use SPE, increasing reward stochasticity would not affect the learning rate, and hence lead to larger updates.

On the neural level, SPE learning predicts trial-by-trial changes of how the dopaminergic prediction error is normalized to a given reward signal. Neural recordings from the relevant brain areas during the learning phase could be compared with simulations of the SPE model to test the theory. In particular, SPE predicts that if reward stochasticity increases slowly, the scale of the corresponding dopamine bursts should stay invariant. Standard theory, on the other hand, would predict that the scale of dopamine bursts grows proportional to the scale of rewards, as prediction errors are a linear function of rewards.

Since the SPE model is closely related to the AU model (11), it inherits a prediction on how the activity in the basal ganglia pathways should depend on reward stochasticity: if reward stochasticity is high, the sum of the activity in the direct and the indirect pathway should be high—if reward stochasticity is low, the sum should be low as well. It has been indeed observed that the neural activity in striatum increases with reward uncertainty (29, 30). Cell-type specific imaging techniques such as photometry could be used to further assert whether the uncertainty is encoded in the sum of activity of striatal neurons on the direct and indirect pathways.

### Relation to models in neuroscience

#### The AU model

The SPE model is closely related to the AU model of Mikhael and Bogacz (11)—both models describe how the basal ganglia pathways track reward uncertainty; they also share the distributed encoding of reward statistics. It is thus not surprising that the learning rules of the two models have similarities. However, the SPE model differs from the AU model in several important aspects.

Of course, the scaled prediction error itself is the key new feature that drives most of the interesting effects we investigated in this work. It is through the scaling of the prediction error that our new model puts its estimate of the reward observation noise to good use. The AU model tracks reward noise as well but does not use its estimate to improve learning performance (or for anything else). In contrast, the SPE model explains not only ***how*** to track *σ*, but also ***why***.

Further, the AU model assumes that there are two separate dopamine signals that modulate activity and plasticity of striatal neurons, namely that the tonic level modulates activity, while the phasic bursts trigger plasticity. However, it has been recently demonstrated that even a brief, burst-like activation of dopaminergic neurons changes the activity levels of striatal neurons (31). Additionally, it has been shown that reward prediction errors modulate the tendency to make risky choices (22), and risk attitudes are known to depend on the balance between the direct and indirect pathways (32, 33). In this paper, we demonstrated that a more realistic assumption, that the dopamine signal encoding prediction error also changes the activity levels in striatum, enables scaling of prediction errors by uncertainty.

#### The Kalman-TD model

SPE learning is not the only model that addresses the scaling of dopamine responses. One recent theory— Kalman-TD—explained those responses, as well as other phenomena such as preconditioning, as a consequence of volatility tracking (5). Kalman-TD applies the Kalman filter method to the computational problem of TD learning: reward prediction in the time domain. The resulting model features vector-valued learning rates that constantly adapt to observations and outcomes. It elegantly describes how covariances between cues and cue-specific uncertainties might modulate learning and can be shown to explain several empirical phenomena. However, the Kalman-TD theory does not address the tracking of observation noise (the theory focuses on process noise). It also does not discuss how prediction error scaling might be implemented. We may thus view it as a complement rather than a competitor to the theory presented above.

#### The reward taxis model

Another model was recently proposed to explain the effects reported by Tobler et al. (10) and other phenomena. The model is called ***reward taxis*** (34), and explains the dopaminergic range adaptation using a logarithm: if both rewards and reward expectations were transformed by a logarithmic function, prediction errors would be given by 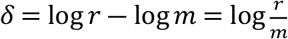. In the experiment of Tobler et al. (10) rewards were given in 50 % of the trials. For a reward of size *r*, the expected reward would then be 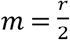, and the prediction error would be 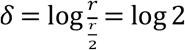, i.e., independent of reward size. Reward taxis can hence explain the results of Tobler et al. (10) quite elegantly.

However, that explanation breaks down as we look at other experiments. We have already mentioned the experiment by Rothenhoefer et al. (21), which featured two reward distributions with equal means and ranges but different standard deviations. We show those distributions in Fig 3C. Rothenhoefer et al. (21) first used Pavlovian conditioning in a way similar way to Tobler et al. (10), pairing the two reward distributions with two different cues. They then recorded the dopamine responses at reward delivery, for all reward sizes of each distribution. We reproduce their data in Fig 3D (first panel). The responses to the middle reward are similar for both distributions, but the responses to the extreme rewards differ: they seem scaled up for the normal distribution.

What would the reward taxis theory predict for the responses in this experiment? Both distributions have the same mean; reward taxis hence predicts similar responses for both distributions. The experimental data thus falsifies the reward taxis model in this experiment. In contrast, the SPE model predicts different responses for the two distributions—we show this in Fig 3D (last panel). Overall, it appears as if dopamine responses to reward distributions with variable width are better captured by the SPE model than the reward taxis model.

#### Free energy models

Finally, we want to discuss the relation of our model to free-energy models: the scaled reward prediction errors in this work are formally related to the precision weighted prediction errors of the free-energy approach, especially when the recognition density (the learner’s model of the world) is taken to be Gaussian (35-37). In that case, the prediction errors that drive inference and learning in free energy models are often weighted by precisions, i.e., inverse variances. The connection to scaled reward prediction errors becomes very close when the free energy approach is applied to reward prediction, dopamine and the basal ganglia system, as has been done in the DopAct framework (38). This framework integrates several theoretical ideas (free energy, reinforcement learning, habits without values and active inference), and suggests that dopaminergic prediction errors drive both learning and action planning.

Precision weighted prediction errors encoded by dopamine transients feature in one variant of that model, but they are not the focus of the theory, and possible implementations or empirical consequences of these weighted prediction errors have not been investigated so far. Furthermore, it is important to note that precision, or inverse variance, scales differently to standard deviation, and might hence not explain classical observations such as those reported by Tobler et al. (10).

### Relation to models in artificial intelligence

Scaled reward prediction errors have been explored outside of neuroscience as well: in the field of AI-type reinforcement learning, it was noticed that normalizing reward prediction errors can enable an agent to learn effectively across several different tasks (39). This is consistent with our conclusions: different tasks come with different levels of reward observation noise, and adaptive scaling can normalize performance across tasks without requiring the need for fine-tuning. However, the rules for scaling prediction errors in AI are different from the SPE learning rules and have not been designed with the intention to model learning in biological systems. Further, Hessel et al. (39) have focused on typical benchmark tasks of AI-type reinforcement learning (i.e. Atari games and others), while we have explored the types of tasks that are used in neuroscience and psychology.

Prediction error scaling also occurs at a more basic level of AI, inside the optimization algorithms that are used to improve the parameters of neural networks. A very prominent example is the Adam optimizer (40), which implements a variant of gradient descent in which all updates are normalized using an estimate of the second moment of the gradient distribution. By making gradient descent effective across different gradient magnitudes, adaptive optimizers such as Adam contribute to the spectacular successes of deep learning. This supports the main idea of this work—that scaling prediction errors can be beneficial for learning. However, here we only looked at the scaling of ***reward*** prediction errors. Adam-style optimization in machine learning, as well as free-energy models in computational neuroscience suggest that there might be similar mechanisms for other neural error signals as well. Therefore, scaling of prediction errors may be a fundamental and common mechanism in the brain. While the mechanisms and evidence presented in this work focus on reward prediction errors and the basal ganglia system, it would also be an interesting direction for future work to investigate scaled prediction errors in other systems within the brain.

## Methods

### Reward prediction performance

In Fig 2, we compare the performance of the RW model with the performance of the SPE model. The SPE model was defined by the learning rules given in Eq. 2 – 4. The RW model was defined by

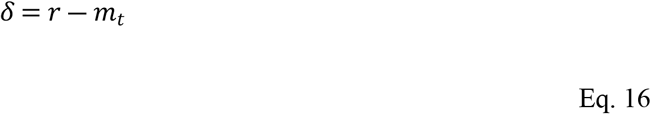

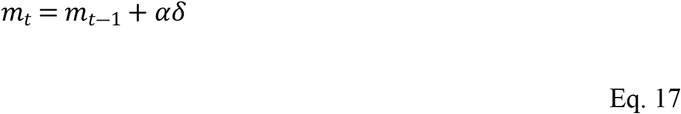

With a constant learning rate *α*.

For Fig 2A, the RW model was parametrized with *α* = 0.5 and the SPE learning rules were parametrized with *α*_*m*_ = 1 and *α*_*s*_ = 0.1. Rewards were sampled from a normal distribution with drifting mean. The process noise was fixed at *v* = 1. We used three different observation noise levels (1, 5 and 15).

For Fig 2B, we used 10 different learning rates for the RW model (ranging from 0.007 to 0.993), the same parameters as above for the SPE model, and 100 different levels of observation noise, evenly distributed on a logarithmic scale from 0.1353 to 1096.6. The process noise was fixed at *v* =1 as above. For each combination, we simulated 10^5^ trials and computed the average squared difference between the model predictions *m* and the true mean *μ* across all trials.

### Simulations of the task of Tobler et al

To simulate the relevant parts of the experiment reported by Tobler et al. (10), we modelled Pavlovian conditioning with three different stimuli, which were associated with three different reward magnitudes (*r* = 0.05, *r* = 0.15, *r* = 0.5). The stimuli were followed by the associated reward in one half of the trials and by no reward in the other half. The rewarded trials were selected pseudorandomly, such that there were two rewarded and two non-rewarded trials in every four successive trials.

We simulated 2000 trials per stimulus and extracted prediction errors from the last 1500. Discarding the first 500 trials accounts for the substantial pretraining of Tobler et al. (10).

We used two models: an RW model and a SPE model. The learning rules of the RW model are given in Eq. 16 and Eq. 17. The rules were used with *α*_*m*_ = 0.0067 and *m*_0_ = 0. The learning rules of the SPE model are given in Eq. 2 – 4. These rules were used with *α*_*m*_ = *α*_*s*_ = 0.0067 and *m*_0_ =0, *s*_0_ = 1.

To compare our simulations to the experimental data from Tobler et al. (10), we extracted the prediction errors *δ* from the simulations and averaged them for each model, outcome and condition separately. There were three conditions (corresponding to the three reward sizes) with two outcomes (reward or no nothing) each, resulting in a total of six combinations per model. Finally, we normalized the six averaged prediction errors by their standard deviation for each model.

### A dynamical model of the basal ganglia

The differential equations Eq. 11 and Eq. 12 were solved using MATLAB’s ***ode15s***, from *t* = − 200 *ms* until *t* = 500 *ms*. As inputs, we used step functions

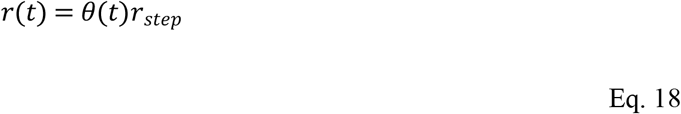

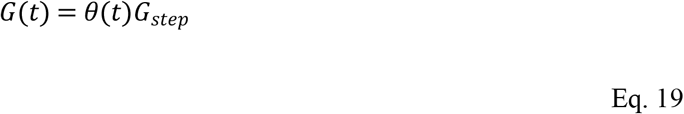

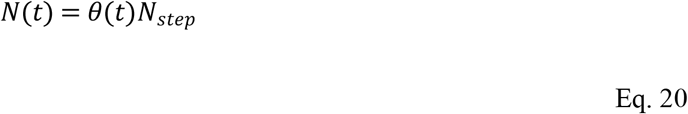

with *θ*(*t*) = 1 for *t* > 0 and *θ*(*t*) = 0 for *t* < 0, and *G*_*step*_ = 10, *N*_*step*_ = 6, and *r*_*step*_ = 4. These inputs correspond to a learned mean *m* = 2 and a learned standard deviation *s* = 8 for *λ* = 1.

The time constant *τ*_*T*_ of the thalamic population was set to 10 ms, based on the measurement of the membrane time constant of thalamic neurons reported by Paz et al. (41). The time constant *τ*_*δ*_ for striatal dopamine was set to 300 ms, based on figure 2C of Montague et al. (42): the dopamine transient in that figure decays to exp −1 of its peak value in about 300 ms.

## Supporting information captions

Appendix S1. Derivation from Bayesian learning

Fig S1. The mode-matching method

Appendix S2. The high noise limit of the steady state Kalman filter

Fig S2. The learning rate of the steady state Kalman filter

## Supporting information

## Appendix S1.

### Derivation from Bayesian learning

One way to derive the scaled prediction error learning rules is the Bayesian mode-matching method, which is a novel (as far as we know) method to approximate Bayesian learning. We first introduce this method. We then apply the method to the problem of tracking the mean and standard deviation of a signal and thus find a new set of learning rules.

#### The mode-matching method

The mode-matching method is based on Bayesian principles. Let us consider the problem of learning the mean and standard deviation of a signal. A fully Bayesian learner would always maintain a belief about the values of the mean and standard deviation, encoded as a probability distribution over all possible pairs of values. It would also maintain a generative model of the signal. When the learner is provided with new information (say another sample of the signal), it applies Bayes’ law to combine its current belief (now the prior) and the likelihood of the observation (computed using the generative model) into a posterior distribution, which encodes its belief after observing the sample. This process is then repeated ad infinitum, with the posterior after one sample turning into the prior for the next.

Now, consider a learner that cannot encode arbitrary belief distributions. Instead, it can only adapt a few of the parameters of a belief distribution with otherwise fixed shape. For example, it might encode a belief using a normal distribution with fixed width and update it by adapting the mean. How might such a learner—let us call it a fixed-shape learner—approximate a fully Bayesian learner best?

Here, we propose the mode-matching method: after observing a new sample, the fixed-shape learner should change the parameters of its belief distribution such that the maximum of the distribution (its mode) is aligned with the maximum of the true posterior. We show this process schematically in Fig S1.

**Fig S1.**
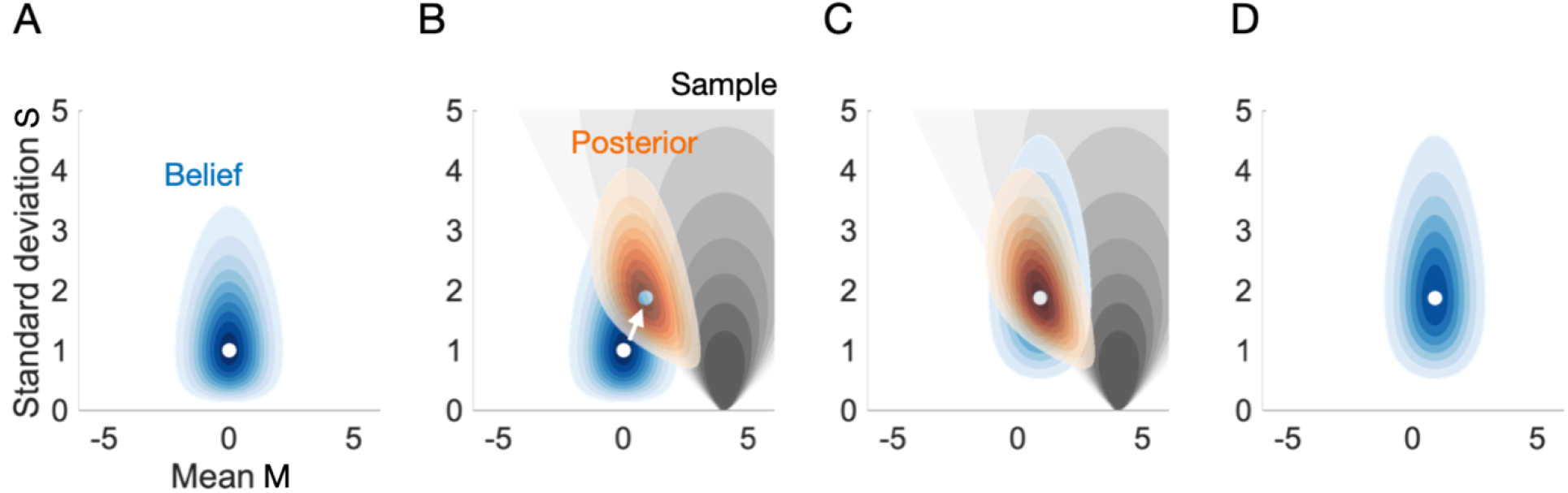
The mode-matching method. The fixed-shape belief distribution is represented by a blue shading; darker shades of blue indicate higher probabilities. Similarly, the true posterior distribution is represented as an orange shading, and the likelihood of the sample is represented by a grey shading. The axes represent the mean and standard deviation of a signal. Units are arbitrary. **A** The fixed-shape belief distribution encodes the learner’s knowledge before a new observation is made. **B** A new observation is made. The likelihood of the observed value is indicated by the grey shading. Using the fixed-shape belief as a prior, a posterior can be computed. The posterior’s mode is different from the mode of the fixed-shape belief; therefore, an update (indicated by a white arrow) is required. **C** The fixed-shape belief has been modified such that its mode aligns with the mode of the true posterior. **D** The modified fixed-shape belief represents the learner’s knowledge after the new observation has been taken into account. The distributions were computed using the densities given in Eq. S7 – S9.

After the update, the fixed-shape learner’s belief is still different from the true posterior. This is because the shape of the true posterior is generally not the same as the fixed belief shape that the learner uses. Hence, mode-matching is only an approximation of Bayesian learning, and some features are lost in this approximation.

Mode-matching is formally related to the variational Bayes scheme (36, 37, 43, 44), which works by minimizing the Kullback-Leibler divergence between the true posterior and a fixed belief shape (usually a multivariate normal distribution). However, mode-matching does not minimise the Kullback-Leibler divergence; instead, it minimises the distance between the modes of the distributions.

The learning rules that can be derived with the mode-matching method are not as precise as those derived from variational Bayes, let alone fully Bayesian learning. What makes mode-matching interesting is that it can be used to derive relatively simple, tractable learning rules, as we shall see in the next section.

#### New learning rules via mode-matching

Let us consider a situation in which an organism tracks the size of a reward associated with some behavior. By engaging in that behavior, it samples the reward size *r*. Using these samples, it attempts to estimate the mean reward *μ* that can be expected from performing the behavior at any given time.

To derive the learning rules for this situation, we start with a generative model for the reward process, and the learner’s fixed-shape belief distributions over the process variables. We model rewards as normally distributed around a mean *μ*, with a standard deviation *σ*, as defined in Eq. 1. Note that *σ* quantifies trial-by-trial fluctuations, and therefore observation noise. The distribution in Eq. 1 is stationary; this means that the environment is modelled as stable.

We further assume that the learner maintains beliefs *M* and *S* about *μ* and *σ*, in form of a normal distribution over possible values of *μ* and a gamma distribution over possible value of *σ*:

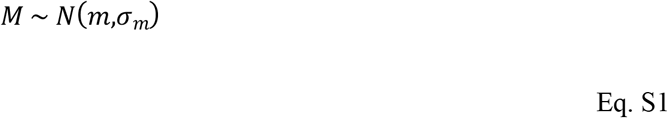

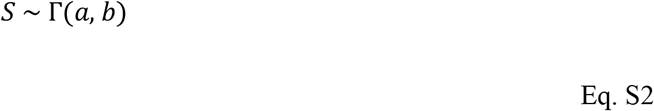

The learner can change its beliefs by adapting the mean (and hence mode) *m* of the normal distribution, and the mode 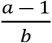 of the gamma distribution. The standard deviation *σ*_*m*_ and the rate parameter *b* stay fixed.

How should we interpret this belief encoding? Allowing *m* and *s* to vary implies that the learner considers both the mean reward *μ* and the observation noise *σ* as unknown—it can adapt its beliefs about these variables. Fixing *σ*_*m*_ and *b* implies that the learner’s uncertainty about the mean and the standard deviation of the signal are kept constant—it cannot adapt those. The learner will thus not become more certain about either the mean or the standard deviation as it gathers more and more data. Fixing *σ*_*m*_ and *b* keeps the resulting learning rules simple. An additional advantage of this design arises when the environment fails to be stationary—then, high certainty about the tracked variables would prevent the learner from adapting to new situations. The model of the reward generating process and the learner’s belief system form our central assumptions—the rest follows. The learner we are about to derive will interpret all rewards it sees as being sampled from a normal distribution with fixed mean and variance, and it will make its inferences accordingly.

Now, let us use mode-matching to derive learning rules from our assumptions. To find out how the learner should update *m* and *s* after sampling a reward *r*, we must first find the mode of the true posterior distribution. For this, we can use a well-known way to simplify calculations. Bayes’ theorem states that

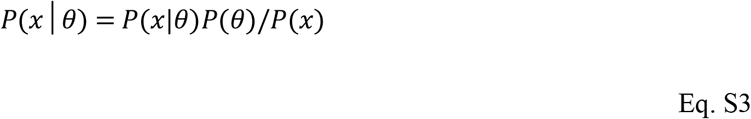

with *θ* the parameters that are to be inferred and *x* the data that is observed. Now we notice that

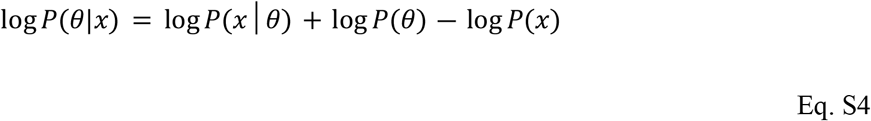

with the last term independent of the parameters θ. We can define the function

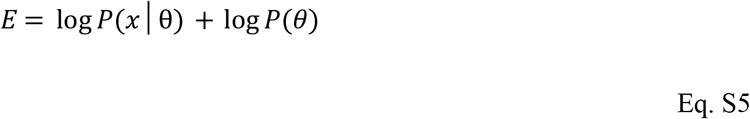

and it is easy to see that the parameters θ_max_ that maximize the function *E* also maximise the posterior distribution *P*(*θ*|*x*) (this is true because the logarithm is strictly monotonic). The function *E*, often called ***energy*** in analogy to statistical physics, is related to the famous of free energy function which plays a key role in many contemporary theories of brain function (38, 45, 46). In the case at hand, the function *E* is given as

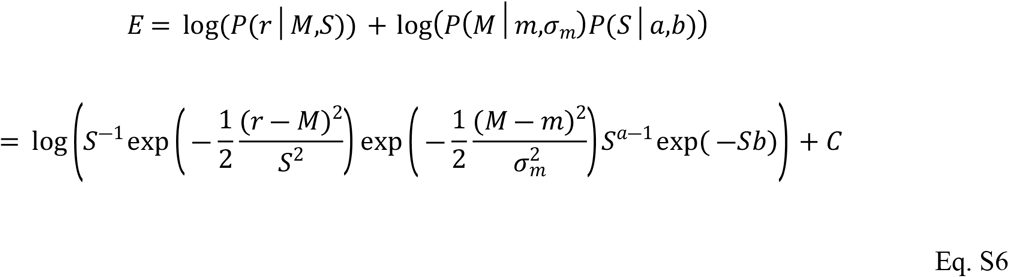

with *C* a term that does not depend on *M* or *S*, and

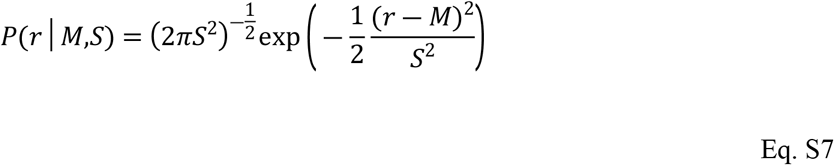

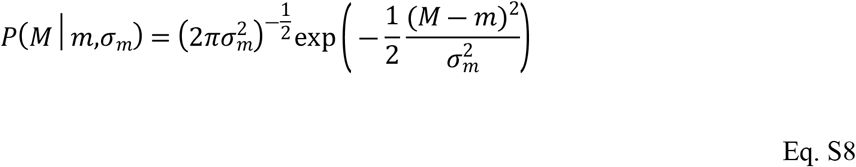

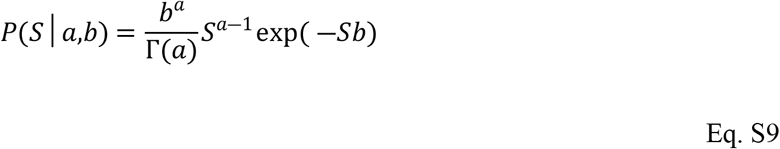

the probability density functions associated with the distributions Eq. 1, Eq. S1 and Eq. S2. To find the maximum of *E* with respect to *M* and *S*, and hence the mode of the posterior, we can investigate the gradient 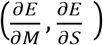 of *E*, which vanishes at the maximum. Evaluation the conditions 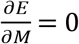 and 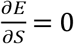, we find

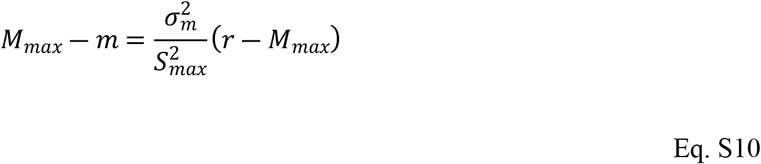

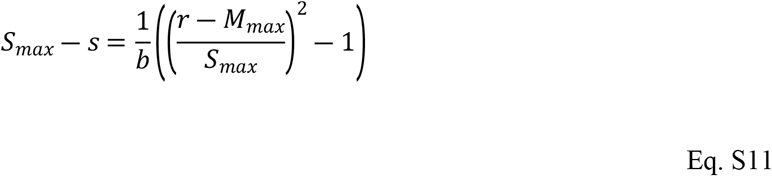

for the location (*M*_*max*_,*S*_*max*_) of the maximum of E. In Eq. S11, 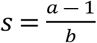 is the mode of the gamma distribution.

To interpret these equations, note that the left-hand side yields the distance of *S*_*max*_ and *M*_*max*_ from the mode of their respective prior distributions. The right-hand side quantifies the mismatch between what was expected based on *M*_*max*_ and *S*_*max*_ and what actually happened: based on *M*_*max*_ and *S*_*max*_, the reward *r* was expected to be close to *M*_*max*_ and (*r* − *M*_*max*_)^2^ was expected to be close to *S*_*max*_ ^2^. The mismatches are weighted with a measure of prior narrowness, 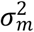 in Eq. S10 and 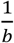 in Eq. S11.

We now must solve these equations for *M*_*max*_ and *S*_*max*_ to find the mode of the true posterior. We could try and find the exact solutions, but considering that the equations are nonlinear, we would have to expect complicated expressions. Here we will not choose that route: we shall restrict ourselves to approximate solutions.

We focus on the scenario in which the priors of both *M*_*max*_ and *S*_*max*_ are very narrow. Formally, this corresponds to 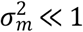 and 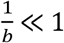, or equivalently 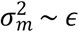 and 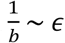 with *ϵ* ≪ 1. To derive an approximate solution for this regime, we use expansions

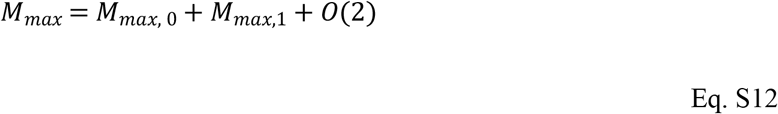

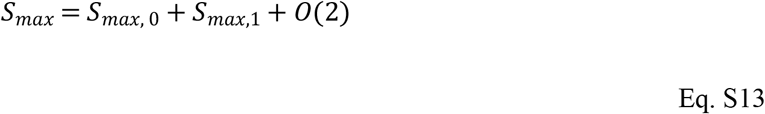

for the variables we want to solve for. Here, *M*_*max*,1_ ∼ *ϵ* and *S*_*max*,1_ ∼ *ϵ* are first order terms with respect to the small constants 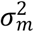 and 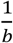. These expansions can be thought of as Taylor expansions of the variables of interest, keeping only terms up to first order. To determine the zeroth and first order terms, we insert these expansions into Eq. S10 and Eq. S11 and collect all terms of a certain order. Using this procedure, we obtain

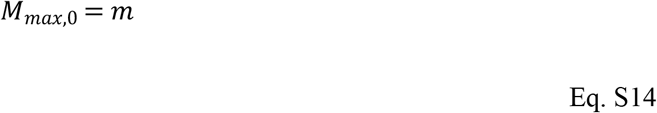

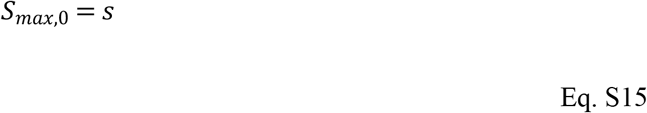

for the zeroth order and

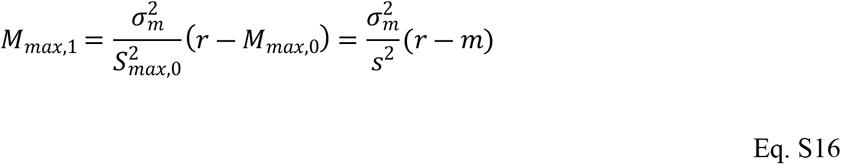

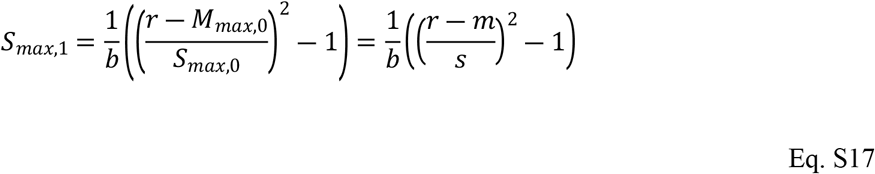

for the first order, where the zeroth order results in Eq. S14 and Eq. S15 were already used. Reinserting these contributions into Eq. S12 and Eq. S13, we find that the mode of the posterior is approximately at

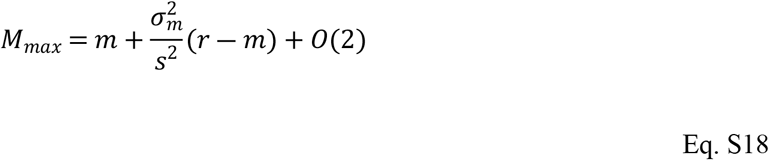

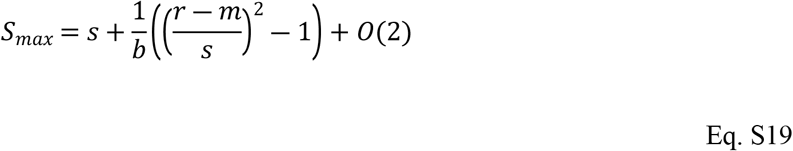

where *O*(2) reminds us that we have neglected terms of second or higher order in 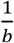 and 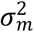. The mode of the posterior is now found—at least approximately. The final step of the mode-matching method consists in updating the mode of the fixed-shape belief distribution—which is (*m, s*)—by aligning it with the maximum of the true posterior, which is (approximately) given by (*M*_*max*_, *S*_*max*_) in Eq. S18 and Eq. S19.

If we were just looking for computationally lightweight learning rules that approximate Bayesian learning, we could stop here. However, we are ultimately interested in modelling learning in biological systems, in particular the basal ganglia system. We must hence consider that changes in synaptic strength can only depend on local information (such as pre- and postsynaptic potentials) and low-dimensional global feedback signals (such as dopamine release in the striatum). We can achieve this here by applying yet another set of approximations. First, we identify certain factors as learning rates: 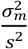 is replaced by *α*_*m*_, and 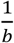 by *α*_*s*_. Then, we simplify the equations by making the learning rates constant; we hence omit the *s*-dependence of *α*_*m*_. With these changes, we arrive at the learning rules specified in Eq. 2 – 4. These rules feature a global feedback signal *δ* and track the mean reward *m* as well as the observation noise *s*. Both *m* and *s* are fed back into the learning system as they enter what we will call the ***scaled*** prediction error *δ*.

## Appendix S2.

### The high noise limit of the steady state Kalman filter

Here, we show that the SPE learning rules approximate the one-dimensional steady-state Kalman filter in the limit of high observation noise. We start by defining the Kalman filter model. We then derive the steady-state Kalman filter, and finally take the high-noise limit.

#### The definition of the Kalman filter

The Kalman filter is a computational method for state estimation and prediction (3). It can be derived from Bayesian principles and is optimal for tracking signals with certain characteristics. Here, we focus on a one-dimensional Kalman filter which is used for predicting rewards, following Piray and Daw (2). The rules they use are

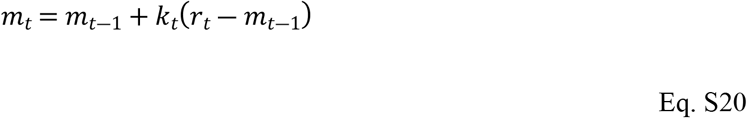

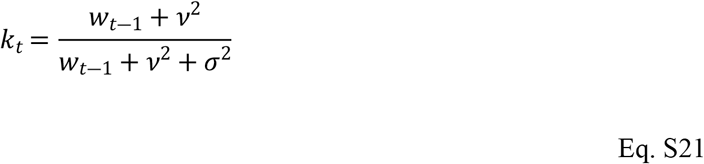

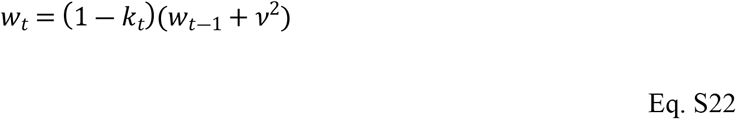

where *r*_*t*_ is the reward, *k*_*t*_ the learning rate or Kalman gain and *w*_*t*_ the posterior variance in trial in trial *t*. Note that our notation differs slightly from that of Piray and Daw (2), for the sake of consistency within this work.

The above rules can be shown to be optimal for tracking signals such as those we used above, i.e., signals that consist of samples drawn from a normal distribution with a drifting mean (3).

#### The steady-state Kalman filter

The Kalman filter has several variables that must be updated on every trial. If one requires a simpler model with almost similar properties, one option is to use a Kalman filter in the limit *t* → ∞: as for *t* → ∞, the posterior variance *w*_*t*_ and the Kalman gain *k*_*t*_ converge to limits *w*_∞_ and *k*_∞_.

Eq. S20 with *k*_∞_ instead of *k*_*t*_ is called a ***steady-state Kalman filter***. By construction, the normal Kalman filter becomes more similar to the steady-state Kalman filter the more trials pass. In practice, performance often does not differ much between the two (3).

What are the limits *w*_∞_ and *k*_∞_? One may use Eq. S21 and Eq. S22 to determine them. By setting *k*_*t*_ = *k*_*t*−1_ and *w*_*t*_ = *w*_*t*−1_, we find

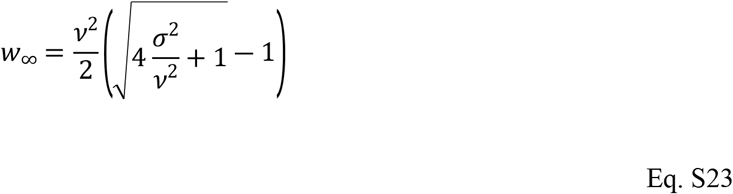

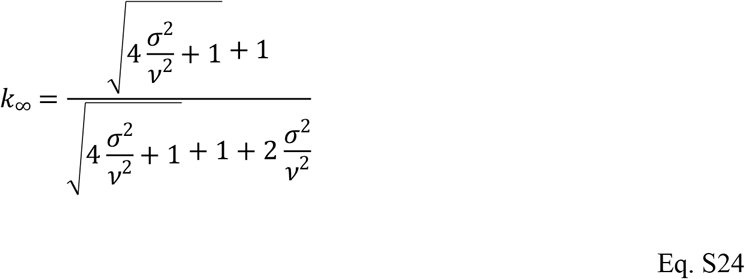

To use the steady-state Kalman filter, one just needs to compute *k*_∞_ and plug it into Eq. S20. One can then use this single equation to track the signal, with no other computations required. The steady-state Kalman filter is thus equivalent to the RW model in Eq. 14 and Eq. 15, parametrized with an optimal learning rate (that is to say, optimal for a signal with statistics *v*^2^ and *σ*^2^).

#### The high-noise limit

The steady-state Kalman filter is less complex than the full Kalman filter. However, its learning rate *k*_∞_ is still a complex function of the signal statistics *v* and *σ*. Can it be simplified? Let us consider of a signal with high observation noise, i.e., with *σ*^2^ much larger than *v*^2^. Using Eq. S24, we find that

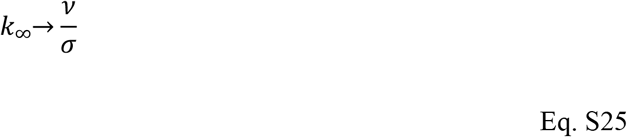

for 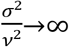. This means that a steady-state Kalman filter with gain 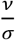 is approximately optimal for signals with *σ*^2^ ≫ *v*^2^. In Fig S2, we compare the optimal steady-state learning rate *k*_∞_ with the approximately optimal learning rate *v*/*σ* for different levels of *σ*, with *v* fixed at *v* =1.

**Fig S2.**
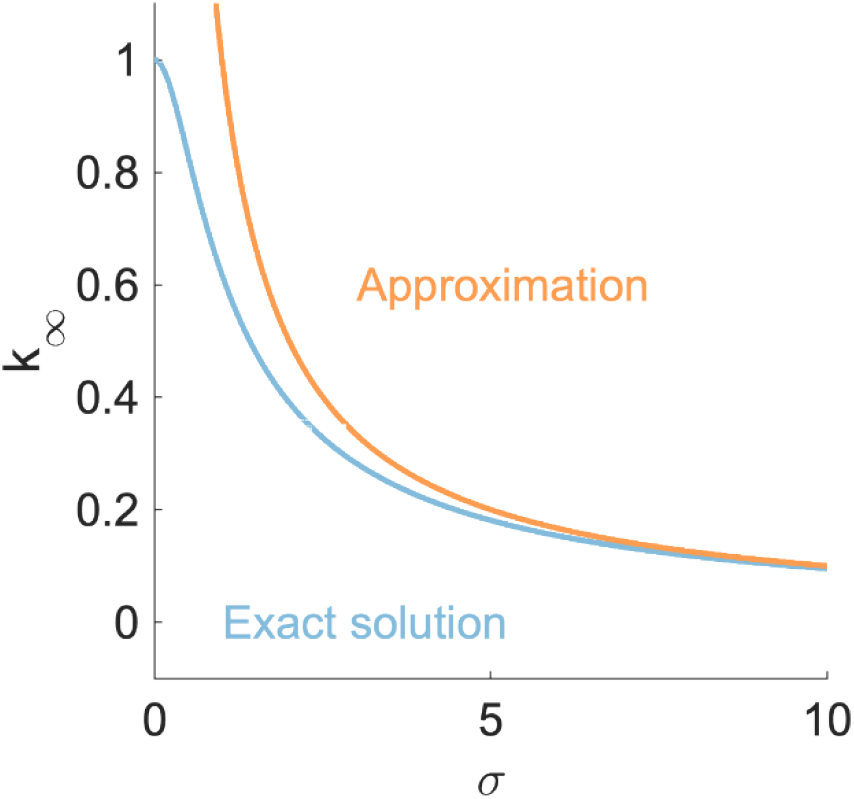
The learning rate of the steady state Kalman filter. We show the learning rate k_∞_ of the steady state Kalman filter as a function of the observation noise σ. We provide the exact value (blue line) and the approximation 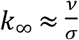 (orange line).

We find that the approximation becomes very close very quickly—for 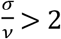, the relative difference between the optimal learning rate and its approximation is already less than 30 %. Fig S2 further suggests that the approximation breaks down as 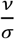 approaches unity—the optimal learning rate for signals with *σ* = 0 is one; any higher learning rate will be detrimental for the performance.

In summary, we find the learning rule

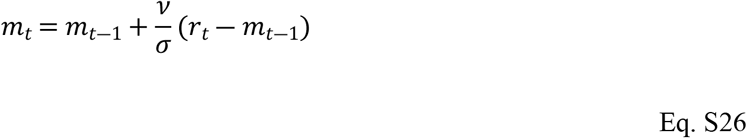

to be approximately optimal for *σ* ≪ *v* and large *t*. The rule Eq. 26 bears striking resemblance to one of the SPE learning rules: the rule in Eq. 3. The difference between the two rules is just how the scaling is attributed: in the Kalman filter, one would perhaps speak of a scaled learning rate, while in the SPE model, one attributes the scaling to the error term. Mathematically, both formulations are equivalent.

A real difference between the Kalman filter and the SPE model is that the latter has a mechanism to track *σ*. No such mechanism exists in the Kalman filter. Both models require *v* as an external input (for the SPE model, the corresponding parameter is *α*_*m*_).

We conclude that the SPE model can be viewed as an implementation of approximately optimal one-dimensional state estimation, equipped with a mechanism to supply some of the required parameters—the observation noise *σ*. Other models have been proposed to track the process noise *v*, for example by Piray and Daw (2). A combination of these approaches might be an interesting direction for future research.

## Notes

### Competing Interest Statement

The authors have declared no competing interest.

